# Omnicrobe, an open-access database of microbial habitats and phenotypes using a comprehensive text mining and data fusion approach

**DOI:** 10.1101/2022.07.21.500958

**Authors:** Sandra Dérozier, Robert Bossy, Louise Deléger, Mouhamadou Ba, Estelle Chaix, Olivier Harlé, Valentin Loux, Hélène Falentin, Claire Nédellec

**Author notes:** Corresponding author, (SD).

## Abstract

The dramatic increase in the amount of microbe descriptions in databases, reports and papers presents a two-fold challenge for accessing the information: integration of heterogeneous data in a standard ontology-based representation and normalization of the textual descriptions by semantic analysis. Recent text mining methods offer powerful ways to extract textual information and generate ontology-based representation.

This paper describes the design of the Omnicrobe application that gathers comprehensive information on habitats, phenotypes and usages of microbes from scientific sources of high interest to the microbiology community. The Omnicrobe database contains around 1 million descriptions of microbe properties that are created by analyzing and combining six information sources of various kinds, i.e. biological resource catalogues, sequence database and scientific literature. The microbe properties are indexed by the Ontobiotope ontology and their taxa are indexed by an extended version of the taxonomy maintained by the National Center for Biotechnology Information.

The Omnicrobe application covers all domains of microbiology. It provides an easy-to-use support in the resolution of scientific questions related to the habitats, phenotypes and uses of microbes through simple and complex ontology-based queries. We illustrate the potential of Omnicrobe with a use case from the food innovation domain.

## Introduction

This paper describes the Omnicrobe database that gathers comprehensive information on habitats, phenotypes and usages of microbes. This information is critical for the development of a large range of research, economic and social activities among which microbial ecosystem services and healthcare are two important topics. Microbial communities render important functions to their ecosystem [1]. These functions are a result of biotic and abiotic interactions dependent on the habitat where microbes live. These last decades, communities have been inventoried by high-throughput culturomics and metagenomic analyses revealing the microbial richness of numerous ecosystems and their ecological importance [2]. Notable examples include the ocean (e.g., Tara Ocean project [3]) and the human gut (Human Microbiome projects [4]). Deciphering microbial traits and phenotypes enables researchers to identify possible beneficial uses for humans in several domains ranging from healthcare or depollution to food [5]. The quantity and the diversity of the information publicly available on focused microbial habitats and phenotypes exponentially increase as a consequence of the growing diversity of microbial ecology research and the parallel evolution of the high-throughput sequencing technologies and novel modeling approaches.

However, methods to assess how microbes are distributed across environments stay limited [6]. The information about microbe habitats and phenotypes is indeed scattered among many different sources, ranging from structured databases (e.g. genetic banks, biological resources, biodiversity databases) to document collections (e.g. scientific literature, reports). Despite the increase in volume and openness of these sources, this wealth of information remains largely underexploited for many reasons: it is distributed in a wide range of sources; the information is described by a great variety of features and metadata that prevents semantic interoperability (e.g. habitat is also called source, isolation, location or host); a large part of the information is expressed in free text even in databases, which makes it difficult to find.

Centralization of microbe habitat and phenotype information by information providers and aggregators (e.g. PubMed Central; https://www.ncbi.nlm.nih.gov/pmc/, GOLD; https://gold.jgi.doe.gov/), standardization of database metadata and increasing use of controlled vocabulary and nomenclatures (e.g. NCBI species taxonomy; https://www.ncbi.nlm.nih.gov/taxonomy) are heading in the right direction for a better access, interoperability and reuse following FAIR principles [7]. The Omnicrobe database project falls fully in line with this effort. Its purpose is to offer a powerful way to retrieve information on microbe taxa, phenotypes, habitats and uses (we define a “use” as any microbial property that can be targeted for human purpose, such as food aromatization) across environments and usages through a single application.

To achieve this goal, the Omnicrobe application automatically aggregates and structures the information that it collects from various public sources (section Information sources). Our work on Omnicrobe focuses on a neglected yet critical aspect: to be easily searched and processed at large scale, the habitat, phenotype and use information gathered on microbes should be organized along standard classifications [8]. Information in Omnicrobe is indexed by a comprehensive controlled and hierarchically organized vocabulary so that the information can be searched in a concise and fast way, regardless of the diversity of terms in source texts. We use text mining methods for parsing and indexing taxon, habitat, phenotype and use mentions and their relationships [9]. The taxa are indexed by the Omnicrobe taxonomic reference, an extended version of the NCBI taxonomy [10,11] and the habitat, phenotype and use information is indexed by the OntoBiotope ontology [12] (section Ontologies and taxonomies). The Omnicrobe database is publicly available at https://omnicrobe.migale.inrae.fr/.

## Background and state of the art

Microbiology covers a wide range of research domains, e.g., molecular biology, ecology, systems biology, evolution, epidemiology, which all deal with microbe environments and phenotypes. The advances in high-throughput technologies generate a tremendous quantity of open access data thanks to the historical investment in shared information systems since the nineties. As a result, data on microbe environments and phenotypes can be found in a diverse range of sources. The most prominent information sources are metagenomic experiment datasets, although the lack of species identification remains an obstacle. The *Genome online database* (GOLD) of the Joint Genome Institute aggregates isolation information from metagenomics experiments on thousands of identified strains [13]. Genetic sequences are also often published with information on isolation samples, as in the GenBank or Biosample databases; this is notably true for complete genome sequences and sequences used for taxon identification (e.g., 16S rRNA gene) [14]. Numerous Biological Resource Centers (BRC) also publish catalogues of their microbial resources with detailed and curated information on the isolation places and phenotypes. The *WFCC Global Catalogue of Microorganisms* aggregates information from 133 collections on almost 500,000 strains with their isolation sources [15]. The *Bacterial Diversity Metadatabase* (*BacDive*) [16] provides information on more than 80,000 strains of the *DSMZ* collection, one of the largest microbe collections in the world. Biodiversity inventories such as *Global Biodiversity Information Facility* (GBIF) and the *Encyclopedia of Life* (EOL) include microorganisms with geographical information but little information on observation places. These information sources can all be searched through open access web pages. The most advanced of them offer an Application Programming Interface (API) access (*e.g. BacDive*). Among these sources, we selected a first core set of databases to populate the Omnicrobe knowledge base, to be extended in the future. These are GenBank, BacDive and INRAE Biological Resource catalogues from the *International Center for Microbial Resources* (CIRM). These sources have been selected for either the quality of their data, their easy access, or their large coverage (section Information sources).

The standardization effort invested in these databases concentrates on taxa information rather than on habitats or phenotypes. The main microorganism references used are *The List of Prokaryotic names with Standing in Nomenclature* (LPSN; https://www.bacterio.net/ [17] (e.g. in *BacDive*)), *the NCBI taxonomy* (e.g. in all NCBI databases and GOLD), and the *Integrated Taxonomic Information System* (in EOL and GBIF). The taxonomic reference of Omnicrobe has a backbone based on the NCBI Taxonomy, with additional strains from BacDive and CIRM. We selected the NCBI taxonomy as microbial taxon reference because it offers a decent coverage of the living, taxa of a wide range of ranks including subspecific ranks and strains, and links to many sequence databases, which opens the way to genomic studies [18].

The information on the microbe isolation sites in databases and articles is expressed in free text fields in various languages. Habitat classification is rarely used to index them. Most habitat classifications used to index microbe isolation sites are non-aligned in-house classifications, notably used by GOLD and BacDive. Isolation sources can actually be searched in BacDive using the *Microbial Isolation Source Ontology* (MISO), a three level-controlled vocabulary of 376 terms [Reimer et al. 2019] that is used to manually index the isolation sources. GOLD classifies the ecosystems of organisms and samples using a five-level controlled vocabulary of 800 terms [13]. The *Earth Microbiome Project Ontology*, EMPO, contains 27 general classes that are relevant for searching correlation patterns between microbial sequences, environment, environmental gradients at a very large scale [6], but not sufficient to record the diversity of the microbial habitats for finer-grained studies. The EnvO ontology offers a larger coverage and a deeper structure for the controlled description of environment types but it is not dedicated to microbes; it is used in projects ranging from plants (e.g. Gramene data resource) to environmental features of some marine species of Tara Oceans expedition [19]. It evolved from the initial objective and is more generally concerned with environments as encountered in ecological applications [20,21]. Its generality and versatility may be an advantage for biodiversity and ecology research at a planetary-scale, but it makes EnvO not as well-adapted to focused domains such as microbiology research.

We chose the OntoBiotope ontology [12] for indexing Omnicrobe habitat information because of its focus on microbe habitats, its richness and its previous uses for text indexing [9]. For phenotype indexing, we also preferred the OntoBiotope classification of microbial phenotypes over other controlled vocabulary, namely the Ontology of Microbial Phenotypes, OMP [22]. OMP focuses on phenotype change (presence, absence, alteration) but lacks some major phenotypes such as morphology phenotypes (e.g. colony color), energy source beyond carbon and oxygen or other environmental factors (e.g. response to various temperature scales).

In the next section, we describe the organization of the database, the controlled vocabulary, the information sources and the text mining process. In the following sections, we present the architecture of the Omnicrobe application, the analysis workflow, the user interface, the content of the current version and the use of Omnicrobe information in the food innovation domain.

## Materials and methods

We present here the schema of the Omnicrobe database, the current data sources and the controlled vocabulary that we use as references to index the data. We also present the text processing method that is applied to index the data sources with the controlled vocabulary.

### Omnicrobe database schema

The Omnicrobe schema is composed of entities of biological interest linked by specific relationships. Four types of entities are defined: *microorganisms, habitats, phenotypes* and *uses*. They are linked by three types of relations: (i) the *lives_in* relation links a microorganism to its habitat; (ii) the *exhibits* relation links a microorganism to its phenotype; and (iii) the *studied_for* relation links a microorganism to its use. This formal schema structures the information in the Omnicrobe database, defines the relevant types of information to be extracted from the data sources and guides the extraction process.

The addition of a new structured source of information requires not only textual information processing of the content but also the alignment of the source schemata with the Omnicrobe data schema. Compared to free-text documents, the relationships between microorganism identifiers and their habitat, phenotype and use descriptions can be more easily extracted from the semi-structured information of the databases. However, the level of information notably varies depending on the databases despite existing efforts on standard schemata for compiling biodiversity data such as the Darwin Core Standard [23] or RDA initiatives (https://rd-alliance.org). Some databases represent the habitats and the geographical locations in separate fields (e.g., Source and Country in GenBank), they may also represent the microbe host or the host part as distinct habitats when relevant (e.g., CIRM CFBP); for the geographical information some databases distinguish the country from the location (e.g., BacDive). It also happens that all this information is mixed in a single field or that the actual content of the database fields does not fully comply with the database schemata, mixing languages for instance. We have thus aligned each information source schema with the Omnicrobe data schema and implemented parsers so that Omnicrobe update is fully automatic.

### Ontologies and taxonomies

The purpose of the Omnicrobe database is data linking and sharing. We have chosen standard controlled reference vocabularies to link microbial data collected from various sources and to make it more findable, accessible, interoperable and reusable, following the FAIR principles.

The OntoBiotope ontology and the NCBI taxonomy have been selected for their relevance to the microbial domain, their lexical richness that makes text parsing more efficient, and their deep structure. Indeed, the deep hierarchical organization of Omnicrobe data enables both smoother browsing by non-specialist users and microbial distribution analysis at various scales.

#### Habitats, Phenotypes and Uses in OntoBiotope

The OntoBiotope ontology (http://agroportal.lirmm.fr/ontologies/ONTOBIOTOPE) [12] contains 4,219 classes split in three branches, Habitat, Phenotype and Use (Table 1). The Habitat branch covers all kinds of microbial isolation sources divided in 11 domains, structured along 13 levels reflecting the diversity of the microbial studies. The Phenotype branch mainly covers physiology, morphology, community behavior, and environmental interaction of various kinds of microbes, e.g., bacteria, fungi, algae. The Use branch describes the uses and applications of microorganisms in 8 subtrees, i.e. antimicrobial activity, pathogenic activity, metabolic activity, with a focus on food, food product quality, sensory quality, mixture transformation, and health properties. The food branch of OntoBiotope, inspired by the FoodEx2 product classification [24], has involved a community effort of INRAE microbiologists to extend it with fermented animal and plant products, cheese, cereal and vegetable juices. These products are the subject of a new and very strong interest for research and innovation [25].

**Table 1.** Number of classes per branch in the OntoBiotope ontology.

| Habitat | Phenotype | Use |
| --- | --- | --- |
| 3,731 | 434 | 66 |

#### Microbial taxa

In order to detect and index microbial “Taxon” entities, we selected the NCBI Taxonomy (https://www.ncbi.nlm.nih.gov/taxonomy) as a reference. The NCBI Taxonomy organizes taxa according to the state-of-the-art phylogeny. The NCBI Taxonomy consists of a classification of taxa in a taxonomic tree, and a nomenclature including valid scientific names, synonyms, and vernacular names, making it easier to standardize taxa in publications even if their name has changed. The NCBI Taxonomy is regularly updated according to new requirements of NCBI databases. The NCBI Taxonomy encompasses all taxa of rank *species* or above, however it only partially covers strains.

Our ambition is to gather information about microorganisms, which is a purely phenotypic notion covering organisms that require a microscope to see them. Therefore, there is no single common ancestor to all microorganisms. We selected a set of 23 high-level taxa (listed in S1 Table) that includes predominantly microscopic individuals, unicellular organisms (bacteria, archaea, virus) and pluricellular organisms (e.g. fungi, algae, nematodes). Omnicrobe is indexed with all sub-taxa of this selection that contains more than 730,000 species. In order to increase the strain coverage in Omnicrobe, we expand the NCBI Taxonomy with strains from the DSMZ catalogue that are publicly available through the BacDive service.

To link each BacDive strain entry to an NCBI Taxonomy node, we automatically match the species and strain names provided by BacDive to taxon names in the NCBI Taxonomy. The objective is not only to gather more strains, but also to place them correctly in the taxonomy and to record synonyms of strain names. The matching process considers several common variations of strain names. The matching has three outcomes:

if one of the names of the BacDive entry or one of its variations is equal to an NCBI Taxonomy strain name, then we add new synonyms to the NCBI strain, and consider the identifiers of the NCBI Taxonomy strain and of the BacDive entry to be equivalent;
if the BacDive entry does not match any NCBI Taxonomy strain, but we can identify the species or genus that the strain belongs to, then we add a new node to the taxonomy;
if the BacDive entry does not match any NCBI Taxonomy strain, nor can we identify which species or genus it belongs to, then we leave this entry out of the reference.

In the current version, the matching process provides 79,041 additional nodes and 796,919 additional synonyms. The Omnicrobe taxonomic reference includes 1,102,673 nodes and 5,862,677 names, including scientific names, vernacular names, and catalogue names for strains. The matching code is Open Source and publicly available at https://forgemia.inra.fr/omnicrobe/extended-microorganisms-taxonomy.

### Information sources

The Omnicrobe database is designed to aggregate information extracted from various sources. The current version includes information from the bibliographic database PubMed (https://pubmed.ncbi.nlm.nih.gov), from the nucleotide database GenBank (https://www.ncbi.nlm.nih.gov/genbank/) and from four microbial resource center catalogues: BacDive (https://bacdive.dsmz.de) and three catalogues of CIRM (*International Center for Microbial Resources*) on food bacteria, plant pathogens and yeast. CIRM BIA (for food bacteria) is available at at https://collection-cirmbia.fr, CIRM-CFBP (for plant pathogens) at https://cirm-cfbp.fr and CIRM-Levures (for yeasts) at https://cirm-levures.bio-aware.com. As stressed above, we have chosen these sources according to their richness, their quality, their popularity, their open access license and their accessibility. Omnicrobe also records the Qualified Presumption of Safety (QPS) of biological agents added to food as maintained by the EFSA (https://www.efsa.europa.eu/en/topics/topic/qualified-presumption-safety-qps).

The information on microorganisms and habitats is extracted by the Omnicrobe application from all the sources, while the information on phenotypes and uses is extracted from Pubmed only.

#### PubMed

The open access bibliographic database PubMed contains more than 33 million citations and abstracts from the biomedical literature maintained by the NCBI and the NLM. Omnicrobe uses a thematic subcorpus of the PubMed references that mention at least one microbe taxon from the Omnicrobe taxonomic reference (section Ontologies and taxonomies). To identify those references, we use an alignment of the Omnicrobe taxonomic reference and the Organisms [B] subtree of the MeSH (Medical Subject Headings) thesaurus that indexes PubMed references. MeSH is available at https://meshb-prev.nlm.nih.gov/search.

At the time of publication of this paper, the Omnicrobe corpus comprises around 2,870,000 PubMed references.

#### GenBank

The GenBank sequence database maintained by the NCBI is an open access, annotated collection of all publicly available nucleotide sequences and their protein translations.

We feed the Omnicrobe taxon field with the parsed content of the “organism” and “strain” fields and the Omnicrobe habitat field with the “isolation_source” field from GenBank releases. The current release (release 249) contains around 5,100 sequence records in the GenBank flat file format. Among these sequences, we select only 16S gene rRNAs sequences because they are generally used to identify bacterial species [26]. The isolation source of the sample is often described by the optional GenBank isolation_source field [14]. We set to 800 base pairs the minimum sequence size to guarantee the quality of the species identification and we use the NCBI Taxonomy part of the Omnicrobe taxonomic reference to filter the relevant GenBank taxa.

#### *BacDive* DSMZ

BacDive is a web service that gives access to the DSMZ catalogue. In 2022, the DSMZ collection maintains more than 80,000 bacterial and archeal strains, including type strains, and is growing fast. The BacDive API returns detailed metadata for each entry such as taxonomy, morphology, physiology, environment and molecular biology. Most of BacDive data is manually curated. The current version of Omnicrobe covers BacDive taxon and habitat information. We feed the Omnicrobe taxon field with the parsed content of the BacDive fields “Full Scientific Name”, “Strain Designation” and “Culture col. no.”, and the habitat field with “Sample type/isolated from” information. BacDive also offers rich information on phenotypes that we plan to process in future work.

#### CIRM

The *International Center for Microbial Resources* (CIRM), managed by INRAE, preserves more than 15,000 strains of bacteria and yeasts among which a subpart is publicly available. Omnicrobe integrates data from three catalogues: CIRM-BIA, dedicated to bacteria of food interest, CIRM-Levures, dedicated to traditional French ferments and yeasts involved in biotechnologies, and CIRM-CFBP on plant-associated bacteria.

Alignments of CIRM database fields to Omnicrobe fields are given in S2 Table. The upcoming availability of CIRM information through the Microbial Resource Research Infrastructure (MIRRI; https://catalog.mirri.org/page/Strains_catalog_query) will provide a single-point and unified access in the near future.

#### Volume of Omnicrobe sources

Table 2 gives the figures of the current version of Omnicrobe sources, with the last revision dates into brackets. S1 Text gives the queries that are used to gather data from PubMed and GenBank sources.

**Table 2.** Omnicrobe sources, data volumes, types and extraction dates in the May 2022 version.

| Source | Volume and types of data [date] |
| --- | --- |
| PubMed | 2.8 million abstracts from articles concerning microbes [2022-04] |
| GenBank | 492,031 entries extracted from the Genbank database. The entries record information about species, strains, isolation sources and hosts [2022-04] |
| BacDive DSMZ | 28,779 entries recording taxa and habitats [2018-01] |
| CIRM-BIA | 1,726 entries recording taxa, strains and habitats [2022-04] |
| CIRM-Levures | 2,158 entries recording taxa, strains and habitats [2021-08] |
| CIRM-CFBP | 6,988 entries recording taxa, strains and habitats [2022-04] |
| QPS species of the EFSA | 788 bacteria with Qualified Presumption of Safety (QPS) status [2022-01] |

### Text mining process

A good part of the information contained in the above-mentioned sources is unstructured and expressed in natural language, either in scientific abstracts (PubMed) or in free-text fields of the databases (GenBank, BacDive DSMZ, CIRM). The text mining process aims to automatically extract this information and structure it according to the Omnicrobe data schema in order to populate the Omnicrobe database.

The text-mining process relies on Natural Language Processing (NLP) techniques and combines rules, lexical resources, and linguistic analysis to detect relevant information from text. The NLP methods themselves are detailed in an earlier work [27] and we will only describe here the general process. It consists of three main steps, namely entity recognition, entity normalization and relation extraction. The aim of entity recognition is to detect textual expressions (or terms) that are of interest for a specific application domain (here, these correspond to microorganisms, habitats, phenotypes, and uses). Then, these entities are normalized with reference knowledge resources (taxonomies, ontologies). That is, each textual entity is linked to a specific entry in a given resource, with a unique identifier and a corresponding label. Here, entities are normalized according to two knowledge resources: microorganisms are mapped to taxa from the Omnicrobe taxonomic reference, while habitats, phenotypes and uses are mapped to concepts from the OntoBiotope ontology. Finally, the third step links together entities that are in relation, according to the predefined set of relations of the Omnicrobe data schema (lives_in, exhibits and studied_for). Note that in the case of database fields, relations between entities are already known and this third step is skipped. Fig 1 gives an example of the three-step text-mining process.

**Fig 1.**
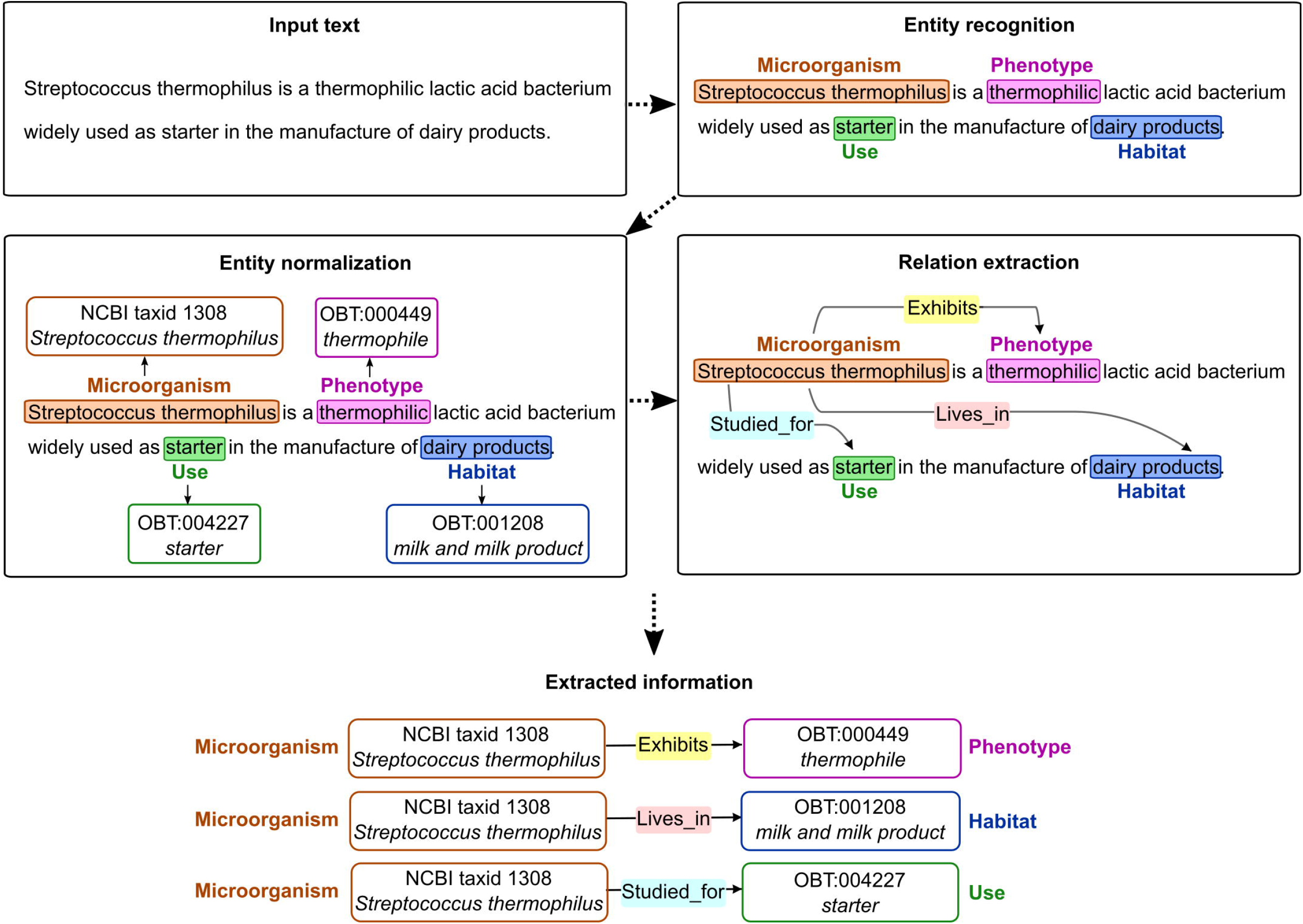
Text-mining process.

The example in Fig 1 is straightforward for illustrative purposes, and does not exemplify the full complexity of extracting textual information. Challenges in text mining stem from the variability and ambiguity of natural language [28]. Automated methods have to deal with numerous linguistic phenomena such as synonymy, abbreviations, homonymy, coreference and complex syntactic structures. Compared to habitats, phenotype and use, taxon naming in scientific texts is more consistent but still a challenge at the strain level, which is highly relevant for many studies [29]. According to [9] respectively 72.5% and 91.2% text mentions do not directly match a category label or synonym of the OntoBiotope classification. S3 Tables give examples of such challenges.

## Results

We designed workflows (section Workflows) to automatically gather, analyze and combine the microbial information in the Omnicrobe Information System (sections User web interface and Application programming interface). Section Omnicrobe content details its current content. The use of Omnicrobe information for food innovation (section Fermentation of soy milk use case) illustrates how relevant data can be retrieved in few queries.

### Information system

As shown in Fig 2, the data from documents is mapped to the Omnicrobe schema and processed by the automatic text-mining workflow. Then this data is integrated into an information system and accessed through a web interface and an application programming interface.

**Fig 2.**
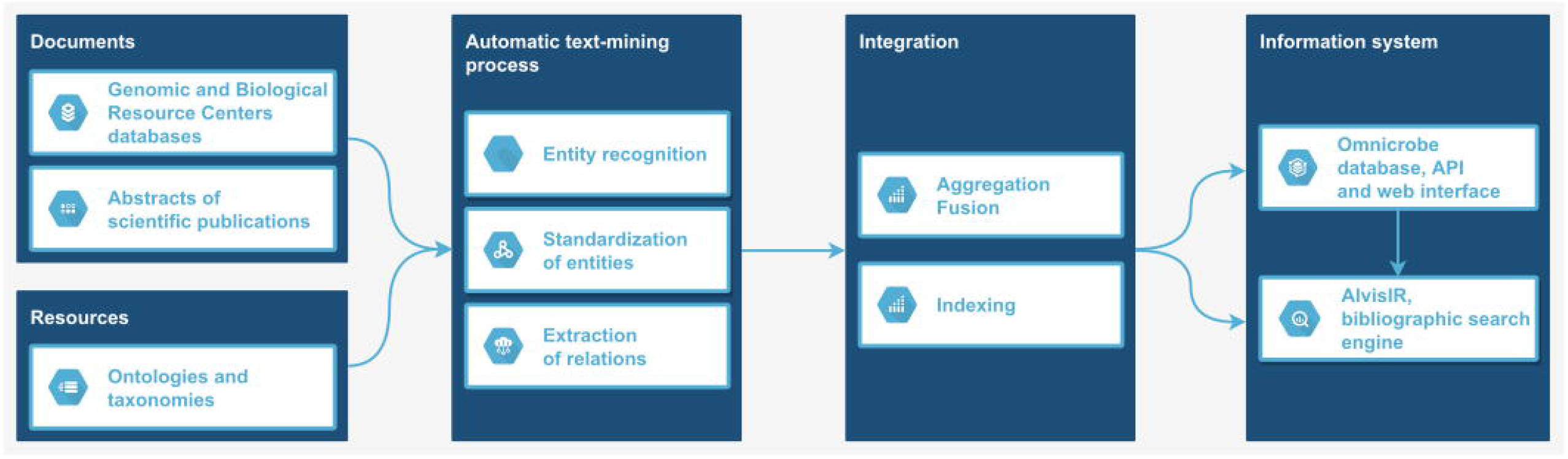
Information system flowchart.

### Workflows

The different processes to generate and integrate data into the Omnicrobe application are implemented into dedicated workflows. These workflows facilitate the management and orchestration of the processing, update and evolution of the Omnicrobe data.

We use Snakemake [30], a workflow management system popular in bioinformatics, to integrate the heterogeneous processing steps and automate the execution. Snakemake manages the tool dependencies and orchestrates the execution of the full workflows on cluster-based environments.

As depicted in Fig 2, several main steps are required to consume the source data and to automatically extract and integrate the results into the information system (i.e. annotations about *microorganisms, habitats, phenotypes* and *uses* and relations between them). These steps are materialized into the following workflows that process and integrate data from each source (https://forgemia.inra.fr/omnicrobe/text-mining-workflow):

**Data collection and preprocessing workflow**: abstracts from PubMed and data from the Genetic and Biological resources centers (GenBank, BacDive and CIRM) are collected and pre-processed by separate workflows. The ontologies and taxonomies are also preprocessed by selecting and filtering the useful parts.
**Text mining workflow**: this workflow implements the text mining process described in the Materials and Methods section (section Text mining process). It ensures the automatic extraction of information from the various data sources (PubMed, GenBank, BacDive and CIRMs). The main text mining steps are implemented using the AlvisNLP text-mining pipeline (https://github.com/Bibliome/alvisnlp). External and third-party tools are used for data pre- and post-processing, i.e., input and output formatting and indexing. The last step of the workflow merges information occurring more than once in the same information source.
**Data integration workflow**: the data integration workflow (https://forgemia.inra.fr/omnicrobe/omnicrobe-database) integrates the data generated by the text-mining workflows into a PostgreSQL relational database. The database indexes the data by the controlled reference vocabulary through ‘hierarchical paths’ so that each entity is indexed by its specific class, and all more general classes of the controlled reference vocabulary.

The Omnicrobe database is regularly updated in order to continuously improve the quality of the data. We continuously enhance the text mining methods according to feedback on the quality of the results and errors observed. We regularly update the information sources and integrate new changes from ancillary resources (e.g., OntoBiotope, Omnicrobe taxonomic reference) when new versions of these resources are released.

The Omnicrobe database has been updated more than 10 times since 2018. The project initially covered the microorganisms and habitats and was extended to the phenotypes and the uses. The input data collected from the different sources has also been notably enriched since 2018 with new articles from PubMed and new BRC.

### User web interface

The Omnicrobe data can be queried through a public web interface (https://omnicrobe.migale.inrae.fr/). It is designed to help microbiologists find information related to their scientific interests. We consulted microbiologists to choose the navigation tools in order to help them make the most of this interface.

Fig 3 shows the main frames of the Omnicrobe web interface. The tabbed navigation (panel A) makes it quick and easy to select the type of search a user wants. For example, users looking for places where a microbe or a family of microorganisms lives, will use the tab “Taxon lives in habitat”. An advanced search functionality is also available and allows advanced users to perform combined multi-criteria searches, for example searching for specific phenotypes of organisms living in a given habitat.

**Fig 3.**
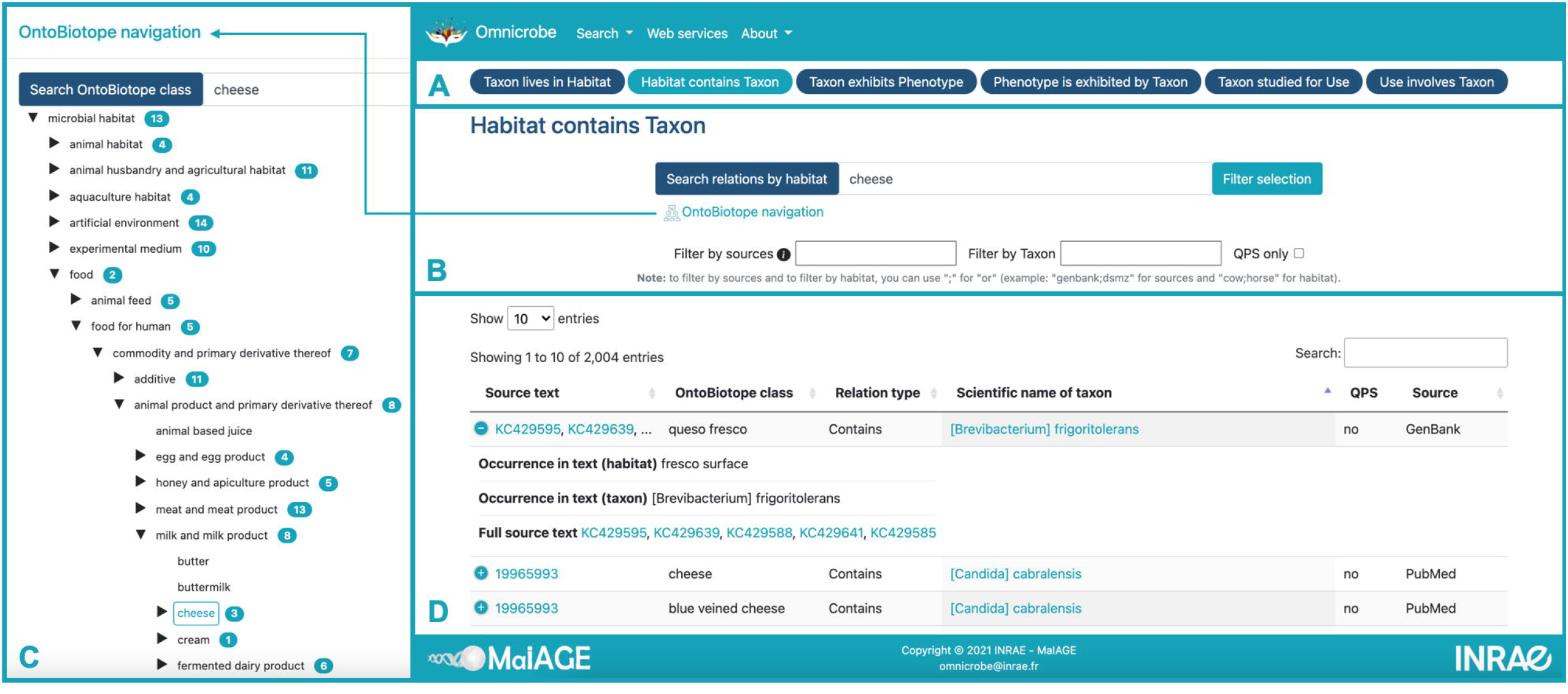
Omnicrobe web interface.

Searches (panel B) are carried out using the terms of habitats, phenotypes and uses of the classes and synonyms of the OntoBiotope ontology and the name of taxon classes of the Omnicrobe taxonomic reference. Queries can be expressed at different levels of generality depending on the needs, from the very specific (e.g., strain, given local specific cheese) to the very general (taxon order, food). In order to optimize searches and avoid any input errors, input fields with automatic completion are proposed. Depending on the type of search, several filters are available, for example, filtering according to the source of the information (e.g., CIRM-BIA) or food safety criteria, i.e., QPS status.

To express his/her query, the user can also navigate in the hierarchical structure of OntoBiotope from the Omnicrobe interface (panel C) and select the class of interest among habitats, phenotypes or uses.

The results (panel D) are displayed in columns for better readability. A link to PubMed and Genbank original text is provided. PubMed references are displayed by the AlvisIR semantic search engine. AlvisIR allows a quick reading of the text thanks to the highlighting of the entities and relations to assess the relevance of the information. Chaix et al. [27] give a detailed presentation of the AlvisIR interface and examples of such use.

The Advanced search of the Search menu is intended for the expression of complex embedded queries that combines “and” and “or” operators. Selected columns of query results obtained from dedicated panels and Advanced search can be exported in various formats (i.e., csv, Excel and pdf) for further processing by users who are not familiar with API.

Omnicrobe runs on an Apache web server. It is written in HTML5/Javascript (client-side) and Python with the Flask web framework (server-side). Using AJAX reduces response time by minimizing data transfers: it sends simultaneous server requests, and takes advantage of the processing capability of the clients.

### Application programming interface

All the data in the Omnicrobe database can be accessed through an Application Programming Interface (API). The API provides the same capability as the web interface. It allows the search for taxa, habitats, and phenotypes, as well as relations between these entities. The main difference is that responses are computer-readable. Thus, the API is suitable to embed Omnicrobe data into another information system. The API is developed with the Python Flask-RESTX framework. The main entry point is available at https://omnicrobe.migale.inrae.fr/api, and the documentation at https://omnicrobe.migale.inrae.fr/api-doc.

### Omnicrobe content

In this section we show an overview of the content of Omnicrobe in its current update (May 2022). The Omnicrobe database content reflects the focus of microbial research studies rather than the worldwide distribution of microbes with respect to their habitat, phenotype or use. The following descriptive statistics reveal the domains where large sets of scientific knowledge has been gathered, and highlight potential gaps that need further attention.

Table 3 shows the number of relations extracted per source after removing duplicates. PubMed is the only source of Omnicrobe for Taxon-Phenotype and Taxon-Use relationships; Integrating Taxon-Phenotype relationships from databases is reserved for future work. PubMed is also the most prolific source for Taxon-Habitat relationships, followed by GenBank. This is expected since PubMed is also the largest source of Omnicrobe.

**Table 3.** Number of distinct relationships extracted per source in the May 2022 version.

| Source | Relationship |  |  |
| --- | --- | --- | --- |
|  | Taxon-Habitat | Taxon-Phenotype | Taxon-Use |
| GenBank | 259,745 | - | - |
| **BacDive** | 35,696 | - | - |
| **CIRM BIA** | 604 | - | - |
| **CIRM Levure** | 943 | - | - |
| **CIRM CFBP** | 1,109 | - | - |
| **PubMed** | 721,244 | 50,410 | 16,007 |

S4 Table lists the most frequent taxa, habitats, phenotypes and uses involved in relationships obtained from PubMed. This ranking reflects the most investigated subjects in papers indexed by PubMed. As expected, they are related to human health: most taxa are pathogens, such as *Staphylococcus aureus*, and a large part of habitats are humans or part of humans, e.g. *patient, blood*, or *respiratory tract*. Phenotypes and uses are related to pathogenicity, resistance, and prevention. The *pathogen* phenotype and *health risk* use are good examples. There is also a sizable focus on fundamental biology (e.g., *cell*, *mouse*). Nevertheless, many other topics are being studied like environmental microorganisms or food contamination (e.g.,*plant, spoilage*).

Fig 4 shows the distribution of taxa ranks. The high number of species and subspecific taxa answers to important needs in microbiology research and is a strong point of the Omnicrobe database.

**Fig 4.**
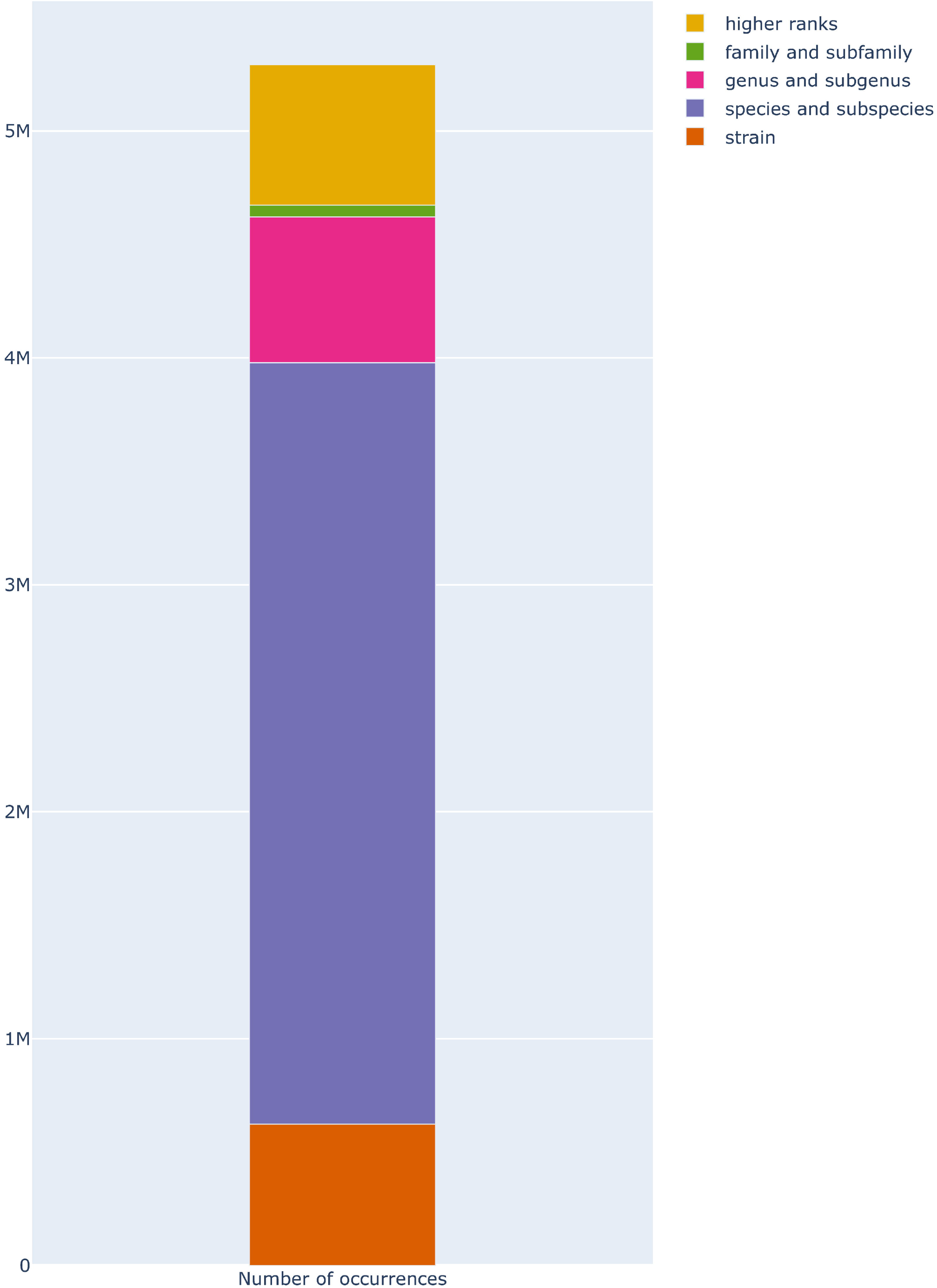
Distribution of taxa ranks in Lives_In relations in Omnicrobe. “Strain” ranks include strains and isolates. “Species and subspecies” ranks include species and ranks below species and above strain (*e.g*., subspecies, varietas, morph). “Genus and subgenus” ranks include genus and ranks below genus and above species (*e.g*., subgenus, section, series). “Family and subfamily” ranks include family and ranks below family and above genus (*e.g*., subfamily, tribe). “higher ranks” include all ranks above the family (*e.g*. order, class, phylum, kingdom). The height of the bars is proportional to the number of *Lives_In* relations in Omnicrobe.

Fig 5 shows the distribution of microbe taxa in *Lives_In* relations. As expected, *Bacteria, Viruses*, and *Fungi* are the most frequent taxa.

**Fig 5.**
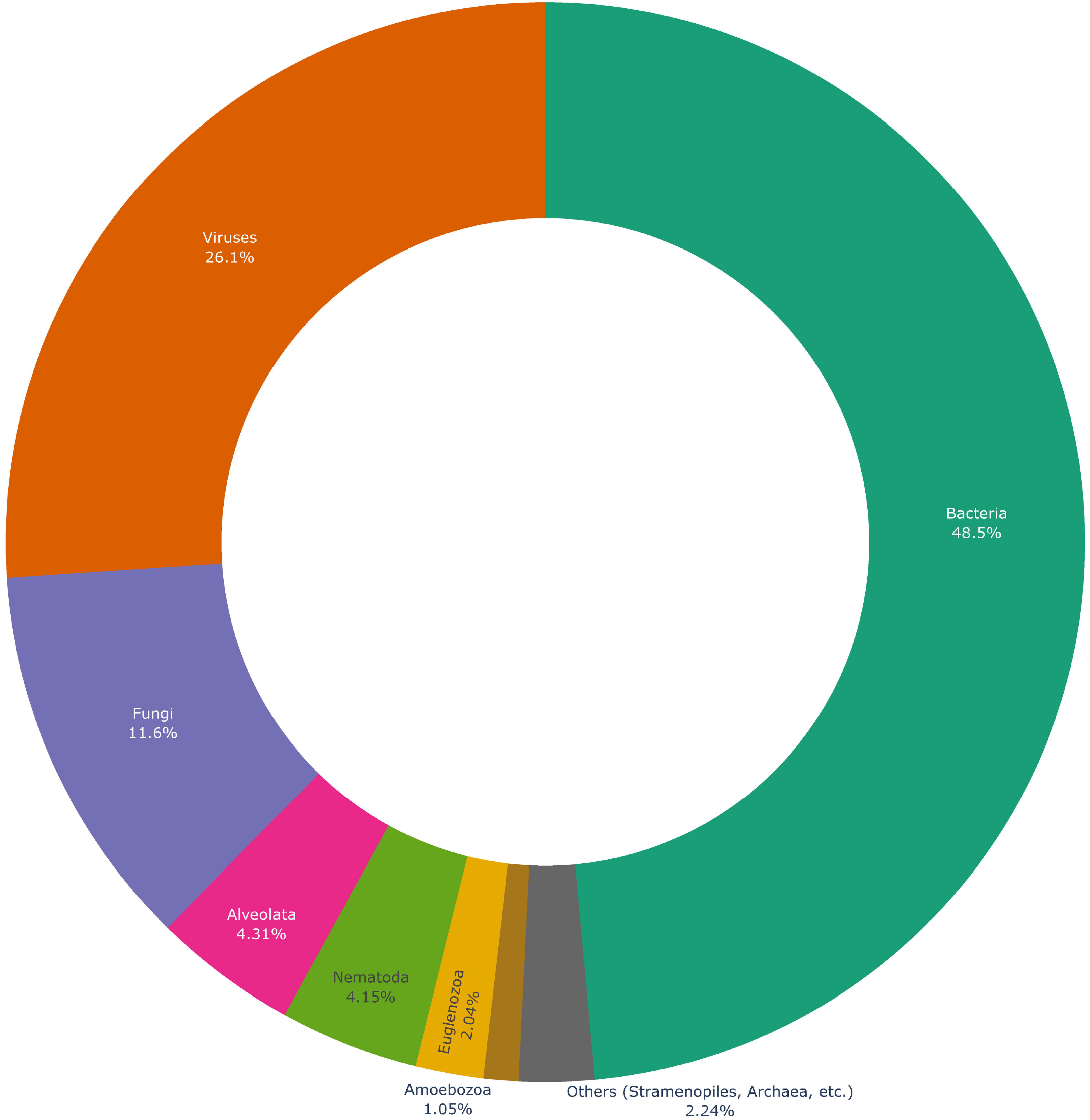
Distribution of microbe taxa in Lives_In relations extracted from PubMed in Omnicrobe. The taxa represented in this chart are taxon roots selected as microorganisms in Omnicrobe (see section Ontologies and taxonomies). The arc is proportional to the number of Lives_In relations that involves the taxon or any descendant. “Others” include taxa that account for less than 1% of relations: *Archae, Chlamydomonadales, Chlorella, Choanoflagellida, Cryptophyta, Desmidiales, Diplomonadida*, Glaucocystophyceae, *Haptophyta, Ichthyosporea, Oxymonadida, Parabasalia, Prototheca, Retortamonadidae, Rhizaria*.

The distribution of habitats at the four highest levels are shown in Fig 6. The most frequent habitats are related to the biomedical domain that provides the major part of Omnicrobe source information. They are followed by engineered (industrial, agricultural, food) environments, non-human hosts and environmental habitats. The distribution is similar to the one shown in the isolation sites distribution chart of BacDive (https://bacdive.dsmz.de/dashboard) [16].

**Fig 6.**
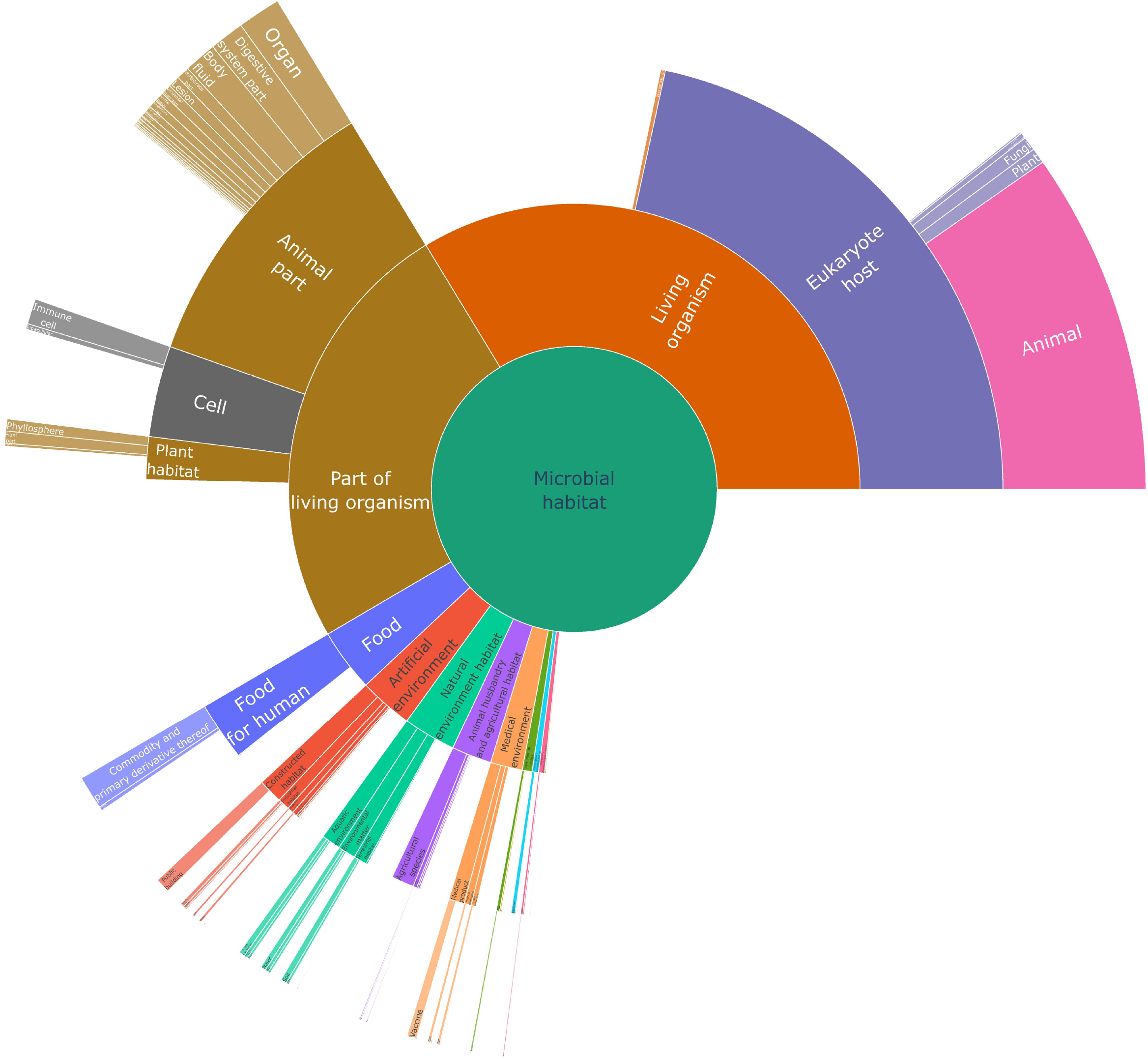
Distribution of habitats in Lives_In relations extracted from PubMed. This chart represents the habitats from the four highest levels in the OntoBiotope ontology. The arc is proportional to the number of *Lives_In* relations extracted from PubMed that involve the habitat or any descendant in OntoBiotope.

Fig 7 shows the proportion of *Lives_In* relations from each source that is also extracted from PubMed abstracts. One might think that the sheer volume of information extracted from PubMed would render the other sources redundant, but this figure indicates the contrary. The overall intersection between PubMed and the other sources is rather low. This demonstrates the complementarity between different sources and that the integration of different sources in Omnicrobe provides more comprehensive information on microbe habitats.

**Fig 7.**
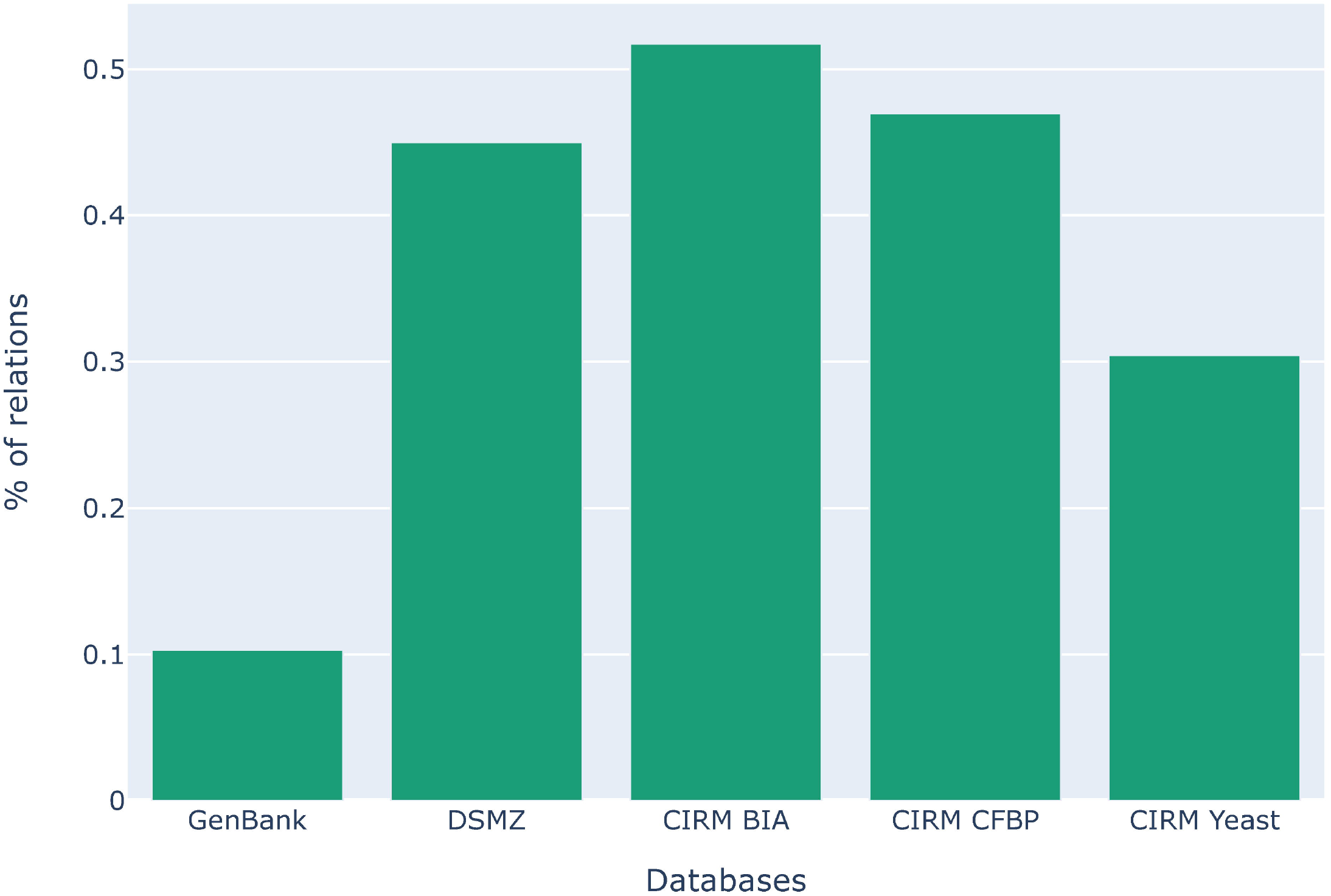
Proportion of taxa-habitat relations in each source that are also extracted from PubMed. The height of the bar represents the proportion of relations per source that were also extracted from PubMed. For instance, only 10% of relations in GenBank were also extracted from PubMed (the same taxon-habitat pair), leaving 90% of relations exclusive to GenBank.

Fig 8 opposes, for the most frequent habitats, the number of relations in which they occur and the number of different microorganism taxa to which they are linked with a relationship. In other words, the frequency of each habitat is contrasted with the diversity of microorganisms that inhabit them. We notice that the diversity is not necessarily correlated to the frequency. For instance, environment and plant habitats (*e.g. plant, yeast, water, soil, marine environment*) display more relative diversity than human and health-related habitats (e.g. human, patient, hospital). Indeed, studies on humans focus on the narrow range of pathogenic microorganisms, whereas studies on plants and environments focus on the biodiversity hosted by these habitats.

**Fig 8.**
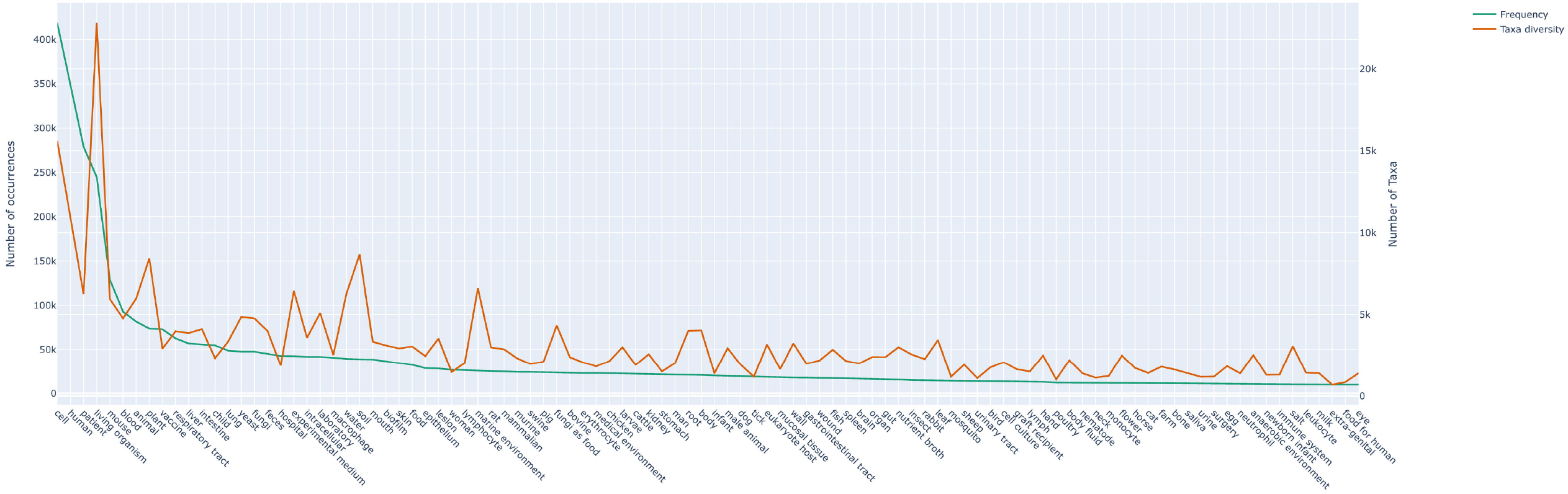
Frequency of habitats and number of different taxa to which they are linked. The green line (left scale) represents the number of *Lives_In* relations extracted from PubMed that involves each of the 100 most frequent habitats. The brown line (right scale) represents the number of distinct taxa to which each habitat is linked with *Lives_In* relations extracted from PubMed.

In order to build an indicator of Omnicrobe consistency, we focused on the six phenotype classes that define the temperature tropism of microorganisms. The classes form a gradient of preferred temperature that ranges from cryophile to extreme thermophile. In this gradient, two phenotypes next to one another (*e.g. thermophile* and *mesophile*) are conceivably compatible. On the other hand, two phenotypes further apart from each other (e.g., *hyperthermophile* and *psychrophile*) are incompatible for the same microorganism. Indeed, it is unlikely that an organism has two very distant optimal growth temperature. Fig. 9 shows the matrix of correlation between all the pairs of the temperature tropism phenotypes. As expected, the diagonal is most intense and the intensity decreases with the distance along the temperature range. This shows that incompatible temperature-based phenotypes are less likely predicted than compatible phenotypes.

**Fig 9.**
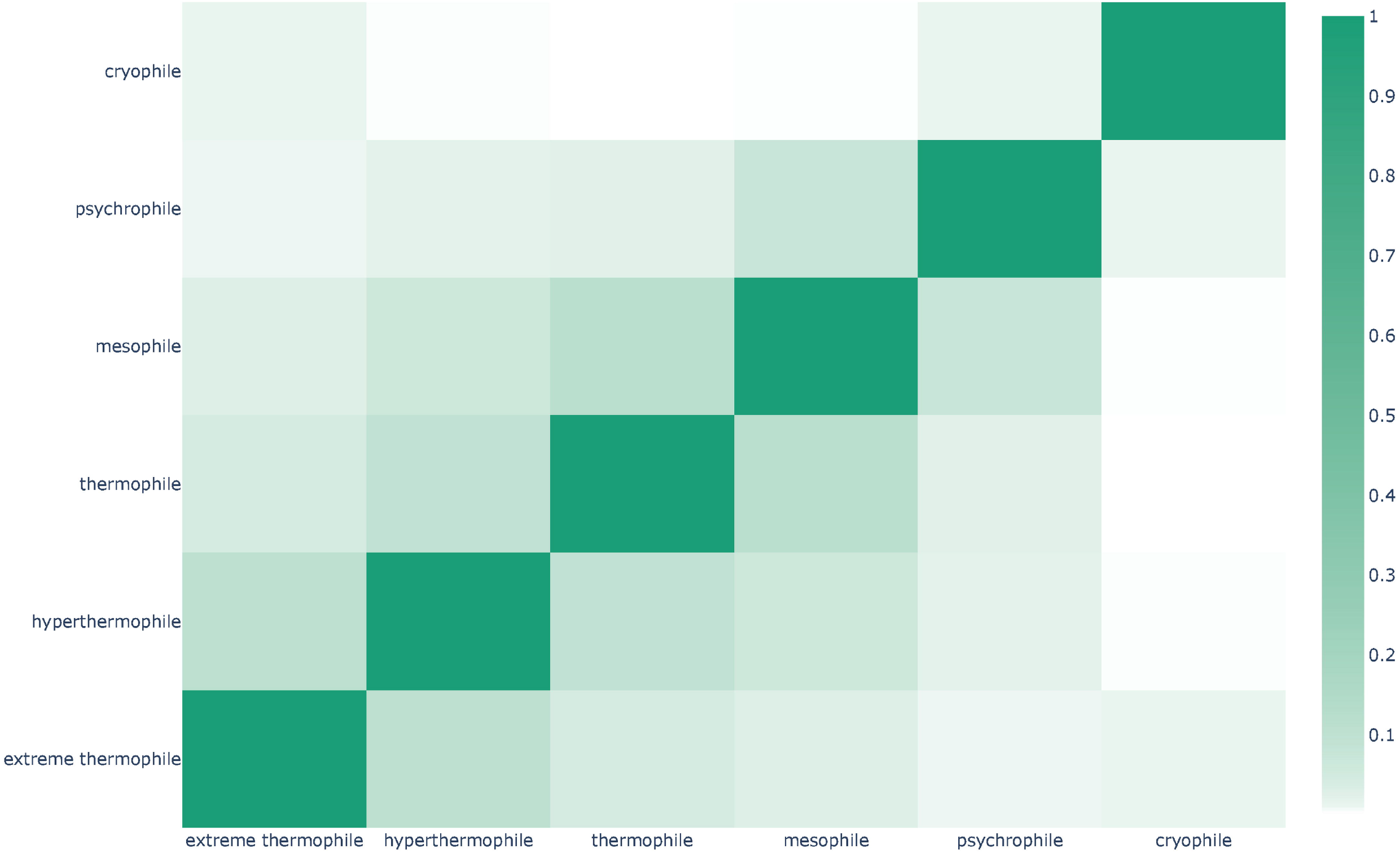
Correlation between temperature tropism phenotypes in Omnicrobe. Each box represents the intersection between the sets of taxa to which the two phenotypes are linked with Exhibits relations in Omnicrobe. The color intensity indicates the Jaccard index between the sets of taxa.

### Fermentation of soy milk use case

Here, we present an example of Omnicrobe application for applied research in food innovation that illustrates both the effective retrieval of relevant information in a short time and its use for further biological development.

#### Use case aim

The aim of this work is to create a new fermented food product based on the fermentation of soy milk [31]. Soymilk represents an interesting alternative to animal milk as a sustainable food. It could also be a valuable protein source for lactose-intolerant and vegan populations. Lactic fermentation of soy juice by lactic acid bacteria (LAB) to produce a yogurt-type fermented soy product can contribute to improving the organoleptic properties of soy juice by reducing “off-flavors” and to lowering the content of non-digestible oligosaccharides. For these reasons, soymilk fermentation attracts recent interest [32].

For this purpose, we would like to identify bacterial species that exhibit the relevant properties. We first search the literature for relevant candidate strains. Once a subset of relevant strains is selected, their *in vitro* cultivation and screening for acidification in soy juice requires the strains to be available in BRC catalogs for ordering. CIRM-BIA dedicated to bacteria of food interest contains more than 4,000 different strains of lactic and propionic acid bacteria and is thus our primary source of strains for this study.

#### Searching Omnicrobe

Classical bibliographic search is complex due to the high number of previous studies and the distribution of the information per bacteria in a high amount of publications, books and websites. We used the Omnicrobe interface to express the combination of criteria that the candidate bacteria have to meet. The targeted properties and their translation into Omnicrobe criteria were defined as follows:

*targeted bacterial species have been previously reported as detected in soy milk* is translated as “soy milk” habitat value
*they are able to perform acidification* is translated as “acidification” use value
*at medium or warm temperature* is translated as “mesophile” or “thermophile” phenotype values
and *they are safe for human food consumption* is translated as QPS criteria set to yes.

The Advanced search of the Search menu was used for the expression of these complex embedded queries by combining “and” and “or” operators as shown in Fig 10.

**Fig 10.**
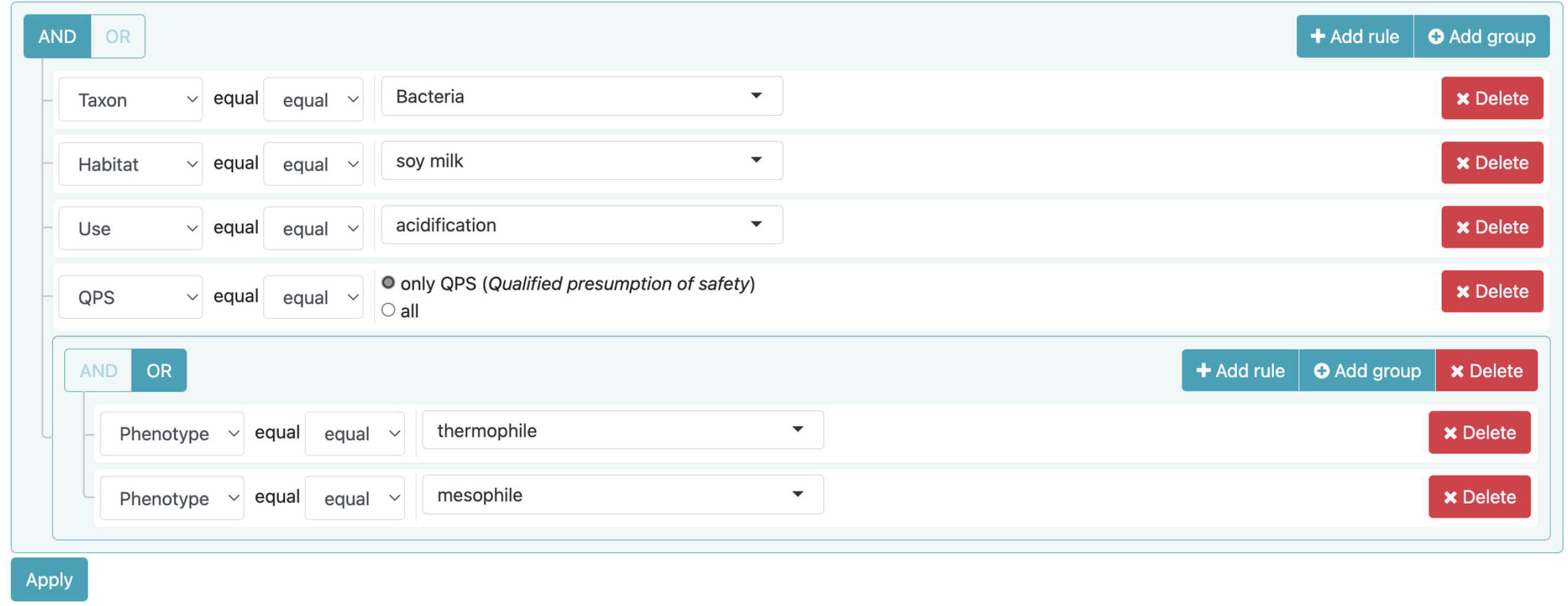
An example of complex embedded queries. Queries are used for the retrieval of mesophilic or thermophilic bacteria present in soy milk and performing acidification and with a qualified presumption of safety.

The resulting list of 103 taxa (20 species, 15 subspecies and 68 strain names) with all requested properties was retrieved from the Omnicrobe interface in a spreadsheet format. The file includes links and IDs of PubMed and GenBank source documents that allow the biologist to check by herself the accuracy of the text mining prediction. The information is spread in 501 documents. For example, *L. acidophilus* has acidification properties according to PubMed ID: 27384493, 9839223, 22264421, 30776138, 9633657, 15591363, and is known to be present in soy milk and soy milk yogurt according to PubMed ID: 18541163, 20477889, 16943081, 28985138, 21775184, 29656125, and is thermophilic according to PubMed ID: 9057296, 1115513.

The analysis of the source texts reveals that two of the retrieved species are actually reported as both thermophile and mesophile: *Lactiplantibacillus plantarum* and *Streptococcus thermophilus*. In the corresponding articles the enumeration of species and phenotypes, and the mixture of species mentions makes difficult the extraction of the relationships by automatic text mining. An example is “A mesophilic-thermophilic mixed culture of *Lactococcus lactis ssp. lactis, Lactococcus lactis ssp. cremoris*, and *Streptococcus thermophilus* was also used.” where *Streptococcus thermophilus* was incorrectly related to mesophilic by the text mining process. We discarded the *Bifidobacterium longum* species because it was considered as ‘obligate anaerobe’ and incompatible with food fermentation conditions.

We aggregated the relationship results at species level. The limited number of species in our study makes manual checking feasible. One additional constraint was the availability of the strains of the selected species in the CIRM-BIA collection (Rennes, France). The eight selected species that meet all requirements were *Lactobacillus acidophilus, Lactobacillus delbrueckii, Lactobacillus helveticus, Lactococcus lactis, Lactiplantibacillus plantarum, Streptococcus thermophilus, Lacticaseibacillus casei, Lacticaseibacillus paracasei*.

#### Fermentation results

We selected 206 strains of the eight species from the CIRM-BIA collection to be tested (S5 Table). The fermentation of the strains was performed in glass vessels with an inoculation of 1 % in the soy juice and a fermentation time of 48 h. Two different temperatures of fermentation were chosen: 30 °C for mesophilic species and 43 °C for thermophilic species. *Streptococcus thermophilus* was cultivated at 43 °C and *L. plantarum* was cultivated at 30°C. The details of the experiment can be found in [31]. Fermentation success was checked by pH monitoring after 48 hours of fermentation. The pH of soy milk was 7.2 at inoculation time. A pH inferior to six means that an acidification process occurred.

Among selected strains, 148 strains succeeded to acidify soy juice in 48 h (S5 Table). All *S. thermophilus* strains acidified soy juice while none of the *L. helveticus* strains acidified soy juice. Acidification is strain-dependent for *L. acidophilus, L. delbrueckii, L. plantarum, L. casei, L. paracasei and L. lactis*. Except for *L. helveticus*, the pre-screening of relevant strains by using the Omnicrobe database was efficient, cost and time effective. The microbiologist saved significant time in bibliographical search and wet lab experiments because she could quickly focus on a relevant subset of species among potential candidates.

## Discussion and conclusion

We presented the Omnicrobe online application, with its unique database of information on microbe habitats and phenotypes. The whole framework was developed to automate as much as possible the update of the database content according to the evolution of the original sources and reference vocabulary and the improvement of the text mining process. In particular, the annotation process of the textual source data and their indexing by standard metadata is fully automatic. Its extension to new available data and sources is straightforward as long as the semantic interoperability of the data is ensured. The increasing standardization of database metadata and use of controlled vocabulary are heading in the right direction. The extension of Omnicrobe with other information available in the current processed sources is the next step. For example, work is underway on Bac*Dive* DSMZ data to extract other types of data than habitat data. Geographical information is one type of data that will require dedicated text mining analysis to ensure semantic interoperability, since the lack of standardization allows mixing in the same record different types of places that should be distinguished, e.g. address, landscape, country. Consideration is also being given to integrating data from GOLD. The scope of the Use part of the database focuses on food application, but will be extended to other domains in future versions, among which biotechnology and sewage treatment.

We presented the Omnicrobe user interface and an example of its use for food fermentation studies. The complementary display by the semantic search engine AlvisIR of the source text and its semantic annotation has proved very useful in checking the quality of the predicted result and putting it into context. The food fermentation use case also confirms the relevance of expressing complex combination of criteria by a query that combines variables and operators. This advanced query feature is intended to users who find the use of the API too technical.

Beyond the inventory of the observations of habitats and phenotypes of microbes offered by Omnicrobe, we think that it could contribute to hypothesize microbe spread scenarios, to anticipate new disease spread and set up adequate control procedures and conversely favor positive flora studies. The ‘One Health’ concept with the interdependence of human, animal, plant and environmental health has stressed the role of new contamination paths that involve intermediate hosts like wildlife, insect vectors, water and air [33]. Microbial ecology of natural environments quickly evolves as a consequence of anthropogenic activities, e.g., climate change, deforestation, water pollution [34], human-wildlife increasing interactions, globalization of plant and animal trade. Predicting the ability of given microorganisms to grow in habitats where they have not been observed strongly depends on the knowledge of their phenotypes and of the connection of their known habitats including vectors and dissemination pathways. Omnicrobe can contribute to provide this information to be used by prediction models such as connectivity models that combines biophysical information, spatial information or air circulation [35].

## Resource availability

Workflows and source code of the Omnicrobe application are distributed on the Git repository (https://forgemia.inra.fr/omnicrobe) under Apache license.

The ontology of microorganism habitats, phenotypes and uses, OntoBiotope, is available on AgroPortal (http://agroportal.lirmm.fr/ontologies/ONTOBIOTOPE).

Information sources: PubMed (https://pubmed.ncbi.nlm.nih.gov), GenBank (https://www.ncbi.nlm.nih.gov/genbank/), BacDive (https://bacdive.dsmz.de), CIRM BIA (https://collection-cirmbia.fr), CIRM-CFBP (https://cirm-cfbp.fr), CIRM-Levures (https://cirm-levures.bio-aware.com) and QPS information from EFSA (https://www.efsa.europa.eu/en/topics/topic/qualified-presumption-safety-qps).

## Acknowledgements

The authors thank the Migale platform for providing the resources to run Omnicrobe services (MIGALE, INRAE, 2020. Migale bioinformatics Facility, doi: 10.15454/1.5572390655343293E12).

The current affiliation of Estelle Chaix is the French Agency for Food, Environmental and Occupational Health & Safety (ANSES), 14 rue Pierre et Marie Curie, 94701 Maisons Alfort Cedex, France.

## Supporting information

**S1 Table. Root taxa of microorganisms included in Omnicrobe.**

**S2 Table. Alignment of CIRM database fields to Omnicrobe database fields.**

**S1 Text. Queries used to extract data from PubMed and GenBank.**

**S3 Table. Examples of challenges faced by text-mining methods.**

**S4 Table. Ten most frequent taxa, habitats, phenotypes and uses in relationships extracted from PubMed in Omnicrobe.**

**S5 Table. Phenotypes registered in Omnicrobe DB V1.0.**

