## Supplementary material for "Omnicrobe, an open-access database of microbial habitats and phenotypes using a comprehensive text mining and data fusion approach": S5 Table

**S5 Table. Phenotypes registered in Omnicrobe DB V1.0**

| **QPS species present in soy juice** | **Species capable of acidification** | **Thermophilic species** | **Mesophilic species** | **Obligate anaerobe** | **Number of *in vitro* tested strain** | **Number of strains leading to acidification (48h)** | **% of strains leading to acidification** |
| --- | --- | --- | --- | --- | --- | --- | --- |
| *Bacillus licheniformis* | X | X |  |  | NA | NA | NA |
| *Bifidobacterium longum* | X | X |  | X | NA | NA | NA |
| *Lacticaseibacillus casei* | X |  | X |  | 5 | 3 | 60 |
| *Lacticaseibacillus paracasei* | X |  | X |  | 9 | 8 | 89 |
| *Lactiplantibacillus plantarum* | X | X | X |  | 44 | 43 | **98** |
| *Lactobacillus acidophilus* | X | X |  |  | 8 | 6 | 75 |
| *Lactobacillus delbrueckii* | X | X |  |  | 21 | 12 | 57 |
| *Lactobacillus helveticus* | X | X |  |  | 16 | 0 | 0 |
| *Lactococcus lactis* | X |  | X |  | 46 | 19 | 41 |
| *Limosilactobacillus fermentum* | X | X |  |  | NA | NA | NA |
| *Streptococcus thermophilus* | X | X | X |  | 57 | 57 | **100** |
| *Total* |  |  |  |  | 206 | 148 | 72 |

NA : fermentation not performed
