## Supplementary material for "Omnicrobe, an open-access database of microbial habitats and phenotypes using a comprehensive text mining and data fusion approach": S4 Table

**S4 Table. Ten most frequent taxa, habitats, phenotypes and uses in relationships extracted from PubMed in Omnicrobe.**

**Table A. Ten most frequent taxa in *Lives_In* relationships**

| **Rank** | **Taxon** | **# *Lives_In* relations** |
| --- | --- | --- |
| **1** | Bacteria | 264360 |
| **2** | *Escherichia coli* | 140159 |
| **3** | Viruses | 126086 |
| **4** | Human immunodeficiency virus 1 | 92914 |
| **5** | *Staphylococcus aureus* | 91717 |
| **6** | Human immunodeficiency virus 3 | 86229 |
| **7** | *Helicobacter pylori* | 85340 |
| **8** | Fungi | 80980 |
| **9** | *Salmonella* | 59983 |
| **10** | Hepacivirus C | 53472 |

**Table B. Ten most frequent taxa in *Exhibits* relationships**

| **Rank** | **Taxon** | **# *Exhibits* relations** |
| --- | --- | --- |
| **1** | Bacteria | 72123 |
| **2** | *Escherichia coli* | 17567 |
| **3** | Fungi | 16142 |
| **4** | Viruses | 7825 |
| **5** | *Staphylococcus aureus* | 7774 |
| **6** | *Pseudomonas aeruginosa* | 5888 |
| **7** | Nematoda | 5615 |
| **8** | *Candida albicans* | 5135 |
| **9** | *Saccharomyces cerevisiae* | 4778 |
| **10** | Archaea | 4047 |

**Table C. Ten most frequent taxa in *Studied_For* relationships**

| **Rank** | **Taxon** | **# *Studied_For* relations** |
| --- | --- | --- |
| **1** | Bacteria | 72123 |
| **2** | *Staphylococcus aureus* | 17567 |
| **3** | *Escherichia coli* | 16142 |
| **4** | *Candida albicans* | 7825 |
| **5** | Fungi | 7774 |
| **6** | *Pseudomonas aeruginosa* | 5888 |
| **7** | *Bacillus subtilis* | 5615 |
| **8** | *Entamoeba coli* | 5135 |
| **9** | *Staphylococcus aureus* DSM 11729 | 4778 |
| **10** | Viruses | 4047 |

**Table D. Ten most frequent habitats in *Lives_In* relationships**

| **Rank** | **Habitat** | **# *Lives_In* relations** |
| --- | --- | --- |
| **1** | cell | 418604 |
| **2** | human | 348892 |
| **3** | patient | 278895 |
| **4** | living organism | 244635 |
| **5** | mouse | 128945 |
| **6** | blood | 92862 |
| **7** | animal | 81564 |
| **8** | plant | 73755 |
| **9** | vaccine | 72805 |
| **10** | respiratory tract | 62593 |

**Table E. Ten most frequent phenotypes in *Exhibits* relationships**

| **Rank** | **Phenotype** | **# *Exhibits* relations** |
| --- | --- | --- |
| **1** | pathogen | 94994 |
| **2** | mutant | 43054 |
| **3** | parasite | 29590 |
| **4** | wild-type | 20799 |
| **5** | anaerobe | 15839 |
| **6** | gram-negative | 13602 |
| **7** | filamentous | 12298 |
| **8** | probiotic | 9603 |
| **9** | phytopathogen | 9386 |
| **10** | aerobe | 9165 |

**Table F. Ten most frequent uses in *Studied_For* relationships**

| **Rank** | **Use** | **# *Studied_For* relations** |
| --- | --- | --- |
| **1** | antibacterial activity | 29463 |
| **2** | antimicrobial activity | 28282 |
| **3** | antifungal activity | 14935 |
| **4** | antiviral activity | 10571 |
| **5** | spoilage | 5576 |
| **6** | starter | 5078 |
| **7** | metabolic activity | 2842 |
| **8** | acidification | 2696 |
| **9** | proteolytic activity | 2660 |
| **10** | health risk | 2455 |
