## Supplementary material for "Omnicrobe, an open-access database of microbial habitats and phenotypes using a comprehensive text mining and data fusion approach": S3 Table

**S3 Table. Examples of challenges faced by text-mining methods.**

**Table A. Challenges when detecting textual entities.**

| **Text with entity mention** | **Ontology concept** | **Challenge(s)** |
| --- | --- | --- |
| *...****commercial soy drink*** *was fermented by 11 lactic acid bacteria...* | Soy beverage | Synonymy |
| *…****soymilk*** *was fermented simultaneously with* Streptococcus thermophilus… | Soy milk | Spelling variation |
| *…supplementation of* ***SM*** *with lactose is likely to enhance the growth of probiotic bacteria...* | Soy milk | Abbreviation |
| *...effects of consuming an* ***isoflavone aglycone-enriched soya milk*** *containing viable bifidobacteria...* | Soy milk | Complex noun phrase and spelling variation |
| *This study evaluated contamination patterns of Listeria species at a poultry food production facility, and evaluated the efficacy of procedures to control the contamination and transfer of the bacteria throughout the* ***plant***. | Food processing factory | Ambiguity of the word “plant” |

The first column shows a text extract with an entity mention, the second column gives the ontology concept corresponding to the textual entity, and the third column explains the type of challenge raised by the example.

**Table B. Challenges when detecting relations.**

| **Text with entity mention** | **Relation between entities** | **Challenge(s)** |
| --- | --- | --- |
| **E. coli *O157*** *was not isolated from any of the 685 export* ***frozen samples*** *tested for this bacteria.* | No relation | Detection of negation |
| *Foodborne disease by* **Listeria monocytogenes, serovar 1/2a** *has recently been reported in many countries. Although contamination by this bacteria is also known to be gradually spreading among the* ***marketed foods of Japan****…* | <Listeria monocytogenes serovar 1/2a> **lives_in** <marketed foods of Japan> | Distinct sentences and coreference phenomenon (“this bacteria”) |
| **Escherichia coli *O157:H7****, an emerging cause of food-borne disease with the occurrence of an estimated 20,000 illnesses and 250 deaths each year in the United States, has now been reported from several countries worldwide. Infections with this bacteria, which follows the ingestion of contaminated food by humans, causes bloody diarrhea, hemolytic uremic syndrome (HUS), and renal disease, that can have serious health implications. The source of food contamination is usually associated with animals, mainly cattle. Many* ***cattle*** *become infected early in life.* | <Escherichia coli O157:H7> **lives_in** <cattle> | Entities are separated by several sentences and the link is implicit |
| *[..] pasteurized milk using a* ***mesophilic*** *lactic acid bacteria culture (***Lactococcus lactis ssp. lactis** *and* **L. lactis ssp. cremoris***) and a* ***thermophilic*** *lactic acid bacteria culture (***Streptococcus salivarius ssp. thermophilus***) [..]* | <Lactococcus lactis ssp. lactis> **exhibits** <mesophilic>  <L. lactis ssp. cremoris> **exhibits** <mesophilic>  <Streptococcus salivarius ssp. thermophilus> **exhibits** <thermophilic> | Multiple candidate couples of entities to relate, which can be difficult to differentiate |

The first column shows a text extract with entity mentions, the second column shows the relation to be extracted between the entities, and the third column explains the type of challenge raised by the example.
