## Supplementary material for "Omnicrobe, an open-access database of microbial habitats and phenotypes using a comprehensive text mining and data fusion approach": S2 Table

**S2 Table. Alignment of CIRM database fields to Omnicrobe database fields.**

| **CIRM** | **CIRM fields** | **Omnicrobe fields** |
| --- | --- | --- |
| CIRM-BIA | Strain_number, Taxon name.Name, ATCC, CIP, DSMZ, LMG, NCIMB, CNRZ, NCDO, NCFB, NCTC CUETM, TL, INA, IL, Other_collection_number  Biotope.Name | Taxon microbien  Habitat |
| CIRM-Levures | CLIB_number, Species name Other_collections  Source_of_isolation | Taxon microbien  Habitat |
| CIRM-CFBP | Especes::Nom_taxonomique, Autres_collections, Autres_noms  Isole_de, Prelevement, Especes::Nom_espece, Especes::Nom_genre, Especes::Nom_sous_espece | Taxon microbien  Habitat (hôte et partie d’hôte) |
