## Supplementary material for "Omnicrobe, an open-access database of microbial habitats and phenotypes using a comprehensive text mining and data fusion approach": S1 Text

**S1 Text. Queries used to extract data from PubMed and GenBank.**

**Text A. MeSH terms used to query PubMed and retrieved references indexed by microbial taxa.**

**https://meshb.nlm.nih.gov**

B01.043

B01.046

B01.050.500.500.294

B01.175

B01.206

B01.237

B01.268

B01.300

B01.400

B01.500

B01.625

B01.630

B01.650.232

B01.650.940.150.469

B01.650.940.150.634

B01.650.940.150.925

B01.650.940.150.950

B01.650.940.800.150.200

B01.675

B01.680

B01.750

B02

B03

B04

**Text B. Genbank data filtering protocol**.

1. Select all sequences matching this expression ‘^.*\\.seq\\.gz’ from GenBank (<https://ftp.ncbi.nih.gov/genbank/>)
2. Filtering sequences: keeping only 16S sequences with a minimal length of 800 bp for bacterial, invertebrates and vertebrates sequences (gbbct*.seq, gbinv*.seq, gbvrl*.seq)
3. Request the following information from GenBank sequences
   a. accession
   b. length
   c. species
   d. strain
   e. taxID
   f. journal
   g. source
   h. host
   i. country
