## Supplementary material for "Omnicrobe, an open-access database of microbial habitats and phenotypes using a comprehensive text mining and data fusion approach": S1 Table

**S1 Table. Root taxa of microorganisms included in Omnicrobe.**

| **Taxon** | **NCBI Taxonomy ID** | **MeSH ID** |
| --- | --- | --- |
| Alveolata | 33630 | D056893 |
| Amoebozoa | 554915 | D056894 |
| Archaea | 2157 | D001105 |
| Bacteria | 2 | D001419 |
| Chlamydomonadales | 3042 | D000077105 |
| Chlorella | 3071 | D002708 |
| Choanoflagellida | 28009 | D056897 |
| Cryptophyta | 3027 | D044785 |
| Desmidiales | 131210 | D058114 |
| Diplomonadida | 5738 | D016828 |
| Euglenozoa | 33682 | D056898 |
| Fungi | 4751 | D005658 |
| Glaucocystophyceae | 38254 | D058108 |
| Haptophyta | 2830 | D058087 |
| Ichthyosporea | 127916 | D050298 |
| Nematoda | 6231 | D009348 |
| Oxymonadida | 66288 | D056899 |
| Parabasalia | 5719 | D056900 |
| Prototheca | 3110 | D011525 |
| Retortamonadidae | 193075 | D056919 |
| Rhizaria | 543769 | D056901 |
| Stramenopiles | 33634 | D058009 |
| Viruses | 10239 | D014780 |
